# Approximate Hidden Semi-Markov Models for Dynamic Connectivity Analysis in Resting-State fMRI

**DOI:** 10.1101/2021.03.01.433385

**Authors:** Mark B. Fiecas, Christian Coffman, Meng Xu, Timothy J. Hendrickson, Bryon A. Mueller, Bonnie Klimes-Dougan, Kathryn R. Cullen

**Affiliations:** Division of Biostatistics, University of Minnesota; University of Minnesota Informatics Institute; Department of Psychiatry and Behavioral Sciences, University of Minnesota; Department of Psychology, University of Minnesota

**Keywords:** Dynamic functional connectivity, fMRI, Hidden Markov models, Hidden semi-Markov models, Multivariate time series

## Abstract

Motivated by a study on adolescent mental health, we conduct a dynamic connectivity analysis using resting-state functional magnetic resonance imaging (fMRI) data. A dynamic connectivity analysis investigates how the interactions between different regions of the brain, represented by the different dimensions of a multivariate time series, change over time. Hidden Markov models (HMMs) and hidden semi-Markov models (HSMMs) are common analytic approaches for conducting dynamic connectivity analyses. However, existing approaches for HSMMs are limited in their ability to incorporate covariate information. In this work, we approximate an HSMM using an HMM for modeling multivariate time series data. The approximate HSMM model allows one to explicitly model dwell-time distributions that are available to HSMMs, while maintaining the theoretical and methodological advances that are available to HMMs. We conducted a simulation study to show the performance of the approximate HSMM relative to other approaches. Finally, we used the approximate HSMM to conduct a dynamic connectivity analysis, where we showed how dwell-time distributions vary across the severity of non-suicidal self-injury (NSSI) in female adolescents, and how those with moderate or severe NSSI have greater state-switching frequency.

## 1 Introduction

The motivating data for this article is the resting-state functional magnetic resonance imaging (fMRI) data obtained from female adolescents in the Brain Imaging Development of Girls’ Emotion and Self (BRIDGES) Study (http://radlab.umn.edu/current-research/bridges-brain-imaging-development-girls-emotion-and-self). For this article, the adolescents were categorized into two clinical groups: adolescents with moderate or severe non-suicidal self-injury (NSSI), and those with no or mild NSSI. The resting-state fMRI data is represented as a multivariate time series, and the time series are independent across the individuals. Our overall objective in this article is to develop a statistical model for *dynamic connectivity analysis*, and determine how dynamic connectivity differs between the two clinical groups.

Dynamic connectivity analysis is the study of how the regions of the brain, which are represented by the different dimensions of the time series, are correlated and how this correlation is potentially changing over time. Dynamic connectivity analysis is still a relatively young approach for fMRI data, but there have been substantial evidence of its utility (Allen et al., 2014; Calhoun et al., 2014; Liégeois et al., 2019; Rashid et al., 2016; Shappell et al., 2021). Existing analytic approaches range from the use of sliding-window correlations (Allen et al., 2014), hidden Markov models (HMM) and hidden semi-Markov models (HSMM) (Vidaurre et al., 2017; Shappell et al., 2019), change-point analysis (Cribben et al., 2012), and time series models with time-varying parameters (Lindquist et al., 2014). Calhoun et al. (2014) gave a thorough overview of analytic approaches for dynamic connectivity analysis.

We focus our attention on HMMs and HSMMs. These models assume that there is a sequence of unobserved states over time, and the properties of the data at each time point within an individual is determined by the current state at that time point for that individual. In our case, we will characterize the states depending on the patterns of *connectivity*, i.e., the structure of the correlation matrix of the multivariate time series. Thus, as the state sequence moves from one state to another state, the correlation matrix will also change. After fitting an HMM or HSMM, one could extract summary statistics such as the number of state switches or the amount of time spent in a state, and one could conduct post-hoc analyses about these summary statistics. HMMs and HSMMs have been useful analytic approaches for modeling multivariate time series data from neuroimaging studies. Indeed, a number of studies have previously used HMMs, HSMMs, or their variants for dynamic connectivity analyses using fMRI data (Baker et al., 2014; Shappell et al., 2019, 2021; Taghia et al., 2017; Vidaurre et al., 2017; Warnick et al., 2018). In addition, neuroimaging studies using other modalities such as electroencephalograms and magnetoencephalograms have also used this modeling framework (Obermaier et al., 2001; Ombao et al., 2018; Quinn et al., 2018). Thus, HMMs and HSMMs have broad appeal for modeling time series data in neuroimaging studies.

HSMMs contain HMMs as a special case. In an HMM, the state sequence is governed by transition probabilities such that the state sequence will either stay in its current state or transition to a different state with a certain probability. Implicit here is that the duration within a state follows a geometric distribution. Consequently, shorter durations occur with higher probabilities. In an HSMM, state transitions are still governed by transition probabilities, but the duration of the state sequence within a state prior to transitioning to a different state is governed by a probability distribution. Thus, an HSMM explicitly models the distribution of the duration within a state, e.g., by assuming that the distribution follows a Poisson distribution or some other discrete distribution. This additional flexibility can affect summary statistics such as the number of times points in a state or how often a state transition occurs. Depending on the context of the problem, practitioners may need the explicit modeling of each state’s duration distribution.

There have been a number of methodological advances for HMMs for multivariate time series. In contrast, the methodological advances for HSMMs are limited in number, especially in the context where we observe multivariate time series data from independent sources, e.g., the female adolescents in the BRIDGES Study. Bulla and Bulla (2006) developed HSMMs for univariate time series data, and software implementation was later provided by Bulla et al. (2010). O’Connell et al. (2011) developed the `mhsmm` package in `R`, which is capable of fitting HSMMs for multivariate time series data from multiple independent sources. Shappell et al. (2019) and Shappell et al. (2021) used this package for the dynamic connectivity analyses in their fMRI studies. An important limitation with their approach is that observed covariates, e.g., NSSI severity, cannot be embedded directly into the model and so the impact of covariates must be investigated in post-hoc analyses. Langrock and Zucchini (2011) showed how one could structure an HMM such that it *approximates* an HSMM and its dwell-time distributions with arbitrary accuracy, and Langrock et al. (2012) illustrated the utility of this approach in modeling animal telemetry data. We build on the modeling framework by Langrock and Zucchini (2011) for multivariate time series data and use it to conduct a dynamic connectivity analysis on the resting-state fMRI time series data from the BRIDGES Study. This modeling framework allows us to have explicit models for the dwell-time distributions that are potentially modulated by covariate information, and we will have available to us the computational tractability and the theoretical and methodological advances already available for HMMs.

The rest of this article is organized as follows. In Section 2, we give a more detailed overview of HMMs and HSMMs, and we describe how one could use an HMM to approximate an HSMM. In Section 3, we use a simulation study to illustrate the performance of the approximate HSMM relative to other models. In Section 4, we show the empirical utility of the approximate HSMM by using it in a dynamic connectivity analysis on the resting-state fMRI data from the BRIDGES Study. Finally, in Section 5 we end with a discussion of the analysis, potential extensions, and limitations.

## 2 Modeling Dynamic Connectivity

### 2.1 Hidden Markov Models and Hidden Semi-Markov Models

We now give an overview of the context of the problem. Let 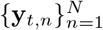 be a collection of *P*-variate time series observed from *N* independent subjects, and we assume that the time series for each subject we have *t* = 1,…, *T*. Denote *y_n_ = (y_1,n_,…, y_T,n_)*. Furthermore, let **Z** be the matrix of covariates whose *n*-th row contains the covariate values for subject *n*. For subject *n*, let *S_t,n_* be an unobserved finite-state process that takes on discrete values in {1,…, *M*} such that, conditional on the states, the observed data ***y**_n_* are independent and follow the *P*-variate normal distribution, also called the *emission distribution*. The mean vectors and covariance matrices of the emission distributions, 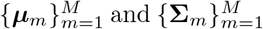, respectively, therefore characterize the states. As the state process switches from one state to the next, the correlation between the dimensions of ***y**_t,n_* also changes. Thus, to model the temporal dynamics of the correlation structure of the time series, our goal is to estimate the covariance matrix in the emission distribution for each of the *M* unobserved states using the collection of both the observed time series 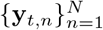 and covariate matrix **Z**.

We now give an overview of the Markov and semi-Markov assumptions for 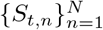 and the hidden Markov and semi-Markov models that arise. If we assume that the *n*-th subject’s finite-state process *S_t,n_* is a Markov process, then for each *n*, we have transition probabilities *a_i,j_* = Pr(*S_t,n_ = j|S_t–1,n_ = i*) with *Σ_j_a_ij_* = 1, and these form the transition probability matrix **A** = (*a_ij_*). Note that the Markov assumption for the state sequence leads to the assumption that the *dwell-time* (or *sojourn time*) within a state, i.e., the number of consecutive time points that the Markov chain spends in a state, follows a geometric distribution. A hidden *semi*-Markov model (HSMM) relaxes this assumption by allowing the dwell-time for state *i* follow a discrete distribution with probability mass function *p_i_(·)* that is potentially parameterized by a vector ***β_i_***. For example, the dwell-time distribution for state *i* could be a Poisson distribution with rate parameter *λ_i_*, and we could have a model for this rate parameter, e.g., log(*λ_i_*) = **Z***β_i_*. The dwell-time distribution characterizes how long the state sequence stays in a state before switching to a different state (i.e., not itself) with some probability. Like the HMM, the state switches in the HSMM are governed by a transition probability matrix **A** = (*a_ij_*), though the diagonal elements of this matrix are 0. The HSMM contains the HMM as a special case if we let each *p_i_(·)* denote the probability mass function for the geometric distribution. We give an illustration of both the HMM and the HSMM in Figure 1.

**Figure 1:**
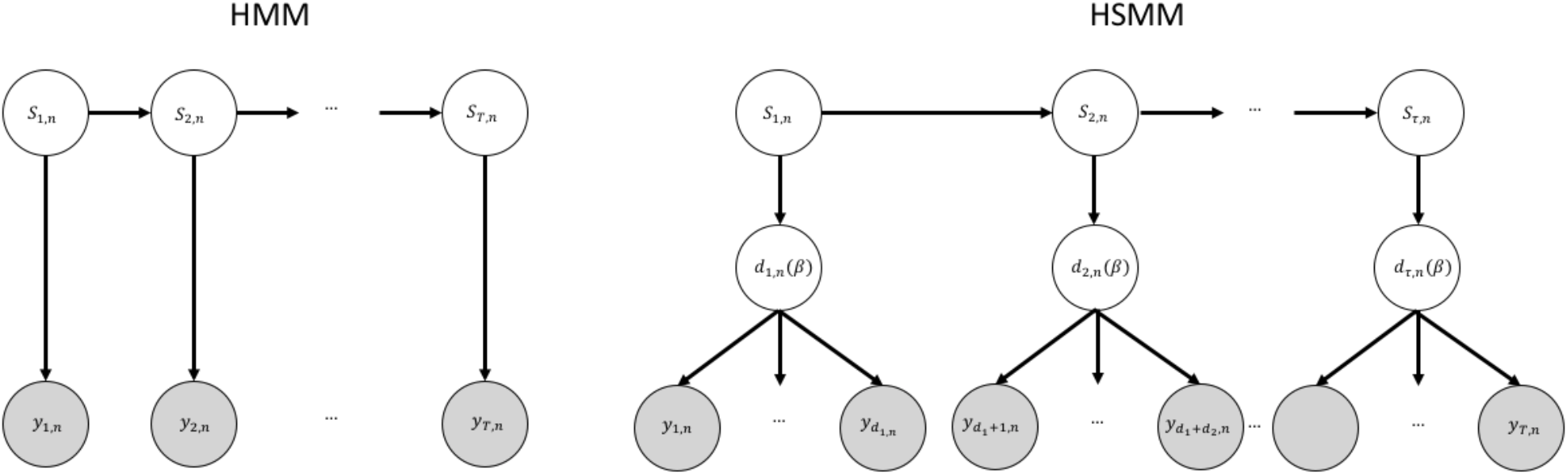
A graphical model representation for a Hidden Markov Model (HMM; left) and a Hidden semi-Markov Model (HSMM; right) for the *n*-th subject. White circles correspond to unobserved variables, and shaded circles correspond to the observed data. For both models, *S_t,n_* is the hidden state sequence that give rise to the observed data **y**_1,n_,…, **y**_T,n_. For the HSMM, *d_1,n_(**β**)*,…, *d_τ,n_(**β**)* are random dwell-times associated with each state that come from a dwell-time distribution with parameter ***β***.

### 2.2 Approximate Hidden Semi-Markov Models

The greater flexibility of the HSMM relative to the HMM results in the likelihood function of the HSMM being more complicated relative to the likelihood function of the HMM, and modeling the dwell-time distribution in the HSMM leads to a substantial computational cost for fitting the model. Our goal is to work in a middle ground in having flexibility in modeling dwell-time distributions with computational complexity. To this end, we follow Langrock and Zucchini (2011) in structuring an HMM in a very deliberate way such that the resulting HMM approximates an HSMM with any form for the dwell-time distributions. In the following, for each *n*, suppose that *S_t,n_* is a *M*-state semi-Markov process with *M x M* transition probability matrix **A** = (*a_ij_*). We now create a Markov process 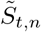 such that an aggregate of its states will approximate *S_t,n_* and its dwell-time. Let *m_1_*,…, *m_M_* be integers with each *m_i_* ≥ 2, and let the state sequence 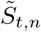 take on values in 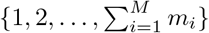. Let **B** be the 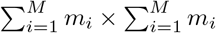 transition probability matrix for 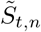, such that it is composed of submatrices, namely,

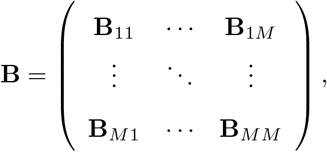

where **B**_ii_ are *m_i_ x m_i_* matrices and **B***_ij_* are *m_i_ x m_j_* matrices. These submatrices have the form

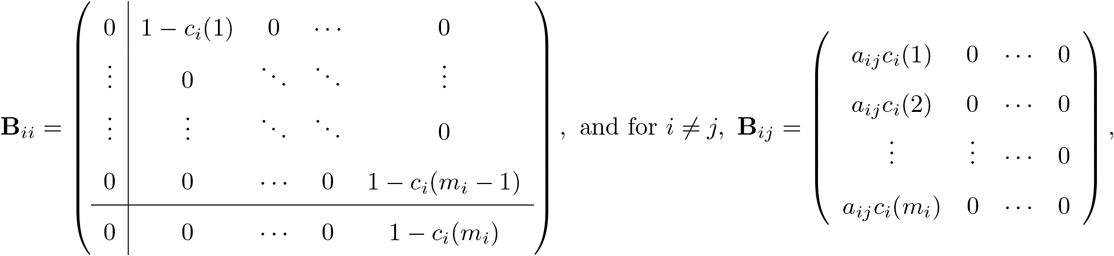

where *c_i_(r)* = *p_i_(r)/(1 – F_i_(r – 1))* for *F_i_(r – 1)* < 1 and *c_i_(r)* = 1 for *F_i_(r – 1)* = 1, and *p_i_(·)* is the probability mass function with cumulative distribution function *F_i_(·)* for the dwell-time distribution of state *i* in the HSMM, with both functions potentially indexed by a parameter vector ***β_i_*** that may be associated with the covariates in **Z**. For *i* = 1,…,*M*, let 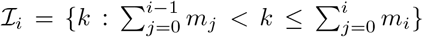, where *m_0_* = 0, be the *i*-th *state aggregate*. 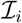 is a collection of states in 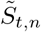 for the approximate HSMM, and is constructed such that 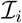 corresponds to state *i* for the state sequences {*S_t,n_*} in the HSMM. Note the deliberate construction of the transition probability matrix **B**. The properties of the *M* states for *S_t,n_* in the HSMM are approximated by the submatrices of **B**. The diagonal block **B***_ii_* represents transitions within state aggregate *i* in the approximate HSMM, and this corresponds to the dwell-time in the HSMM. The off-diagonal block **B***_ij_, i ≠ j*, characterizes the transitions between state aggregates in the approximate HSMM, corresponding to the transitions between states in the HSMM. Note that a transition from state aggregate 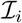 to state aggregate 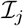 in the approximate HSMM must go to the state with the smallest index in 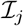.

We now give a few remarks about the approximate HSMM being in an HMM framework, and its relation to the HSMM, which Langrock and Zucchini (2011) discussed in greater detail:

- All entries of **B** lie in the interval [0,1], and its row sums equal to 1, and so **B** is a valid transition probability matrix.
- One can show that the transition from 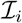 to a different state aggregate 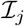 in the approximate HSMM has the same probability of state transition governed by the transition probability matrix **A** in the HSMM.
- Different dwell-time distributions will lead to different formulations for *c_i_(r)*, and thus different entries for each submatrix in **B**. However, the deliberate structure for **B** will remain the same.
- Since the states within 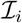 in the approximate HSMM correspond to state i in the HSMM, the parameters of the emission distribution for all states within 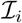 must be constrained to be the same.

The cardinality of 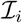 is *m_i_*, and this is a parameter that is set *a priori* for each state aggregate. Langrock and Zucchini (2011) showed that the errors in the approximation is in the right tail of the dwell-time distribution, and that a larger value of *m_i_* leads to a better approximation of the dwell-time distribution. In Figure 2, we show how the choice of *m_i_* affects the approximation of the dwell-time distribution. In the example in the figure, suppose the dwell-time distribution for one of the states in the HSMM is the shifted Poisson distribution with shift parameter 1 and rate parameter *λ*, and suppose the approximate HSMM uses the correctly specified dwell-time distribution. The rows in Figure 2 correspond to dwell-time distributions with different rate parameters, where in the top row we have *λ* = 3 and the bottom row we have *λ* = 10. The three columns in Figure 2 correspond to the different sizes of the state aggregates, namely, *m_i_* = 5, 10, or 15. The vertical lines correspond to the actual values of the dwell-time distribution, and the dots correspond to the values of the dwell-time distribution induced by the approximation that arises for each *m_i_*. In the top row, we see that setting *m_i_* = 10 or 15 approximates the dwell-time distribution really well, but setting *m_i_* = 5 leads to errors in the right tail of the distribution. When *λ* = 3 we see that the error in the approximation when using *m_i_* = 5 is in the right-tail of the distribution, but using *m_i_* = 10 or 15 leads to good approximations. When *λ* = 10 we see that the errors are very pronounced for *m_i_* = 5, but the approximation is much better for larger values of *m_i_*. In fact, we see that the larger the mean for the dwell-time distribution the larger we need to set *m_i_*. Altogether, *m_i_* is a parameter where larger values lead to better approximations, with the trade-off that larger values of *m_i_* will increase the dimensions of **B** and hence will increase the computational cost.

**Figure 2:**
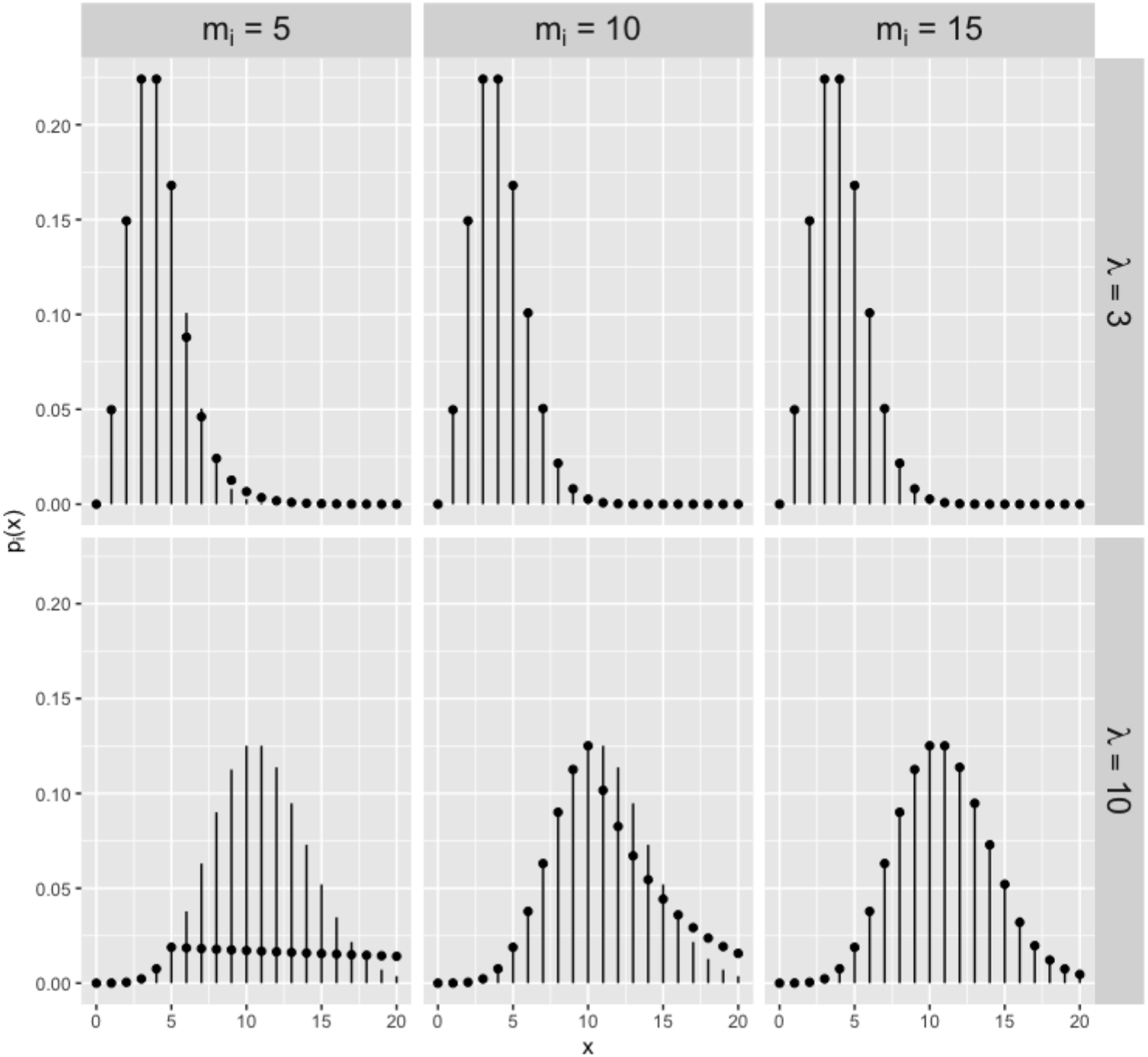
In each plot, we have the shifted Poisson distribution with shift parameter 1 and rate parameter *λ* = 3 (top row) and *λ* =10 (bottom row). The columns correspond to the different sizes *m_i_* of the state aggregates. The vertical lines in each plot shows the probability mass function of the shifted Poisson distribution, and the dots correspond to the induced dwell-time distribution of the approximated HSMM for a given *m_i_*.

**Figure 3:**
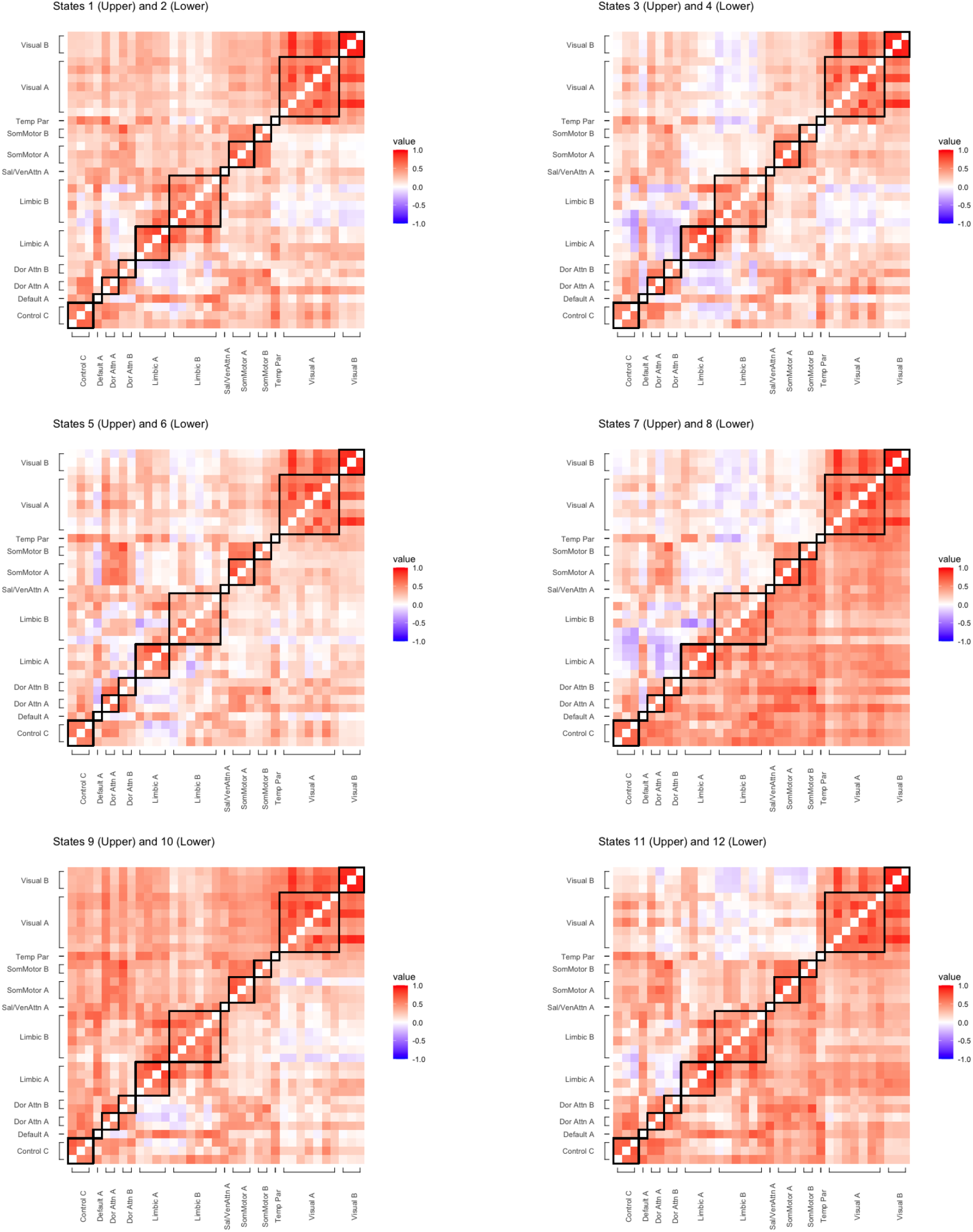
The correlation matrix for each state, with upper and lower triangles corresponding to different states. The black squares within the plot correspond to the ICNs mapped to the same Yeo regions.

**Figure 4:**
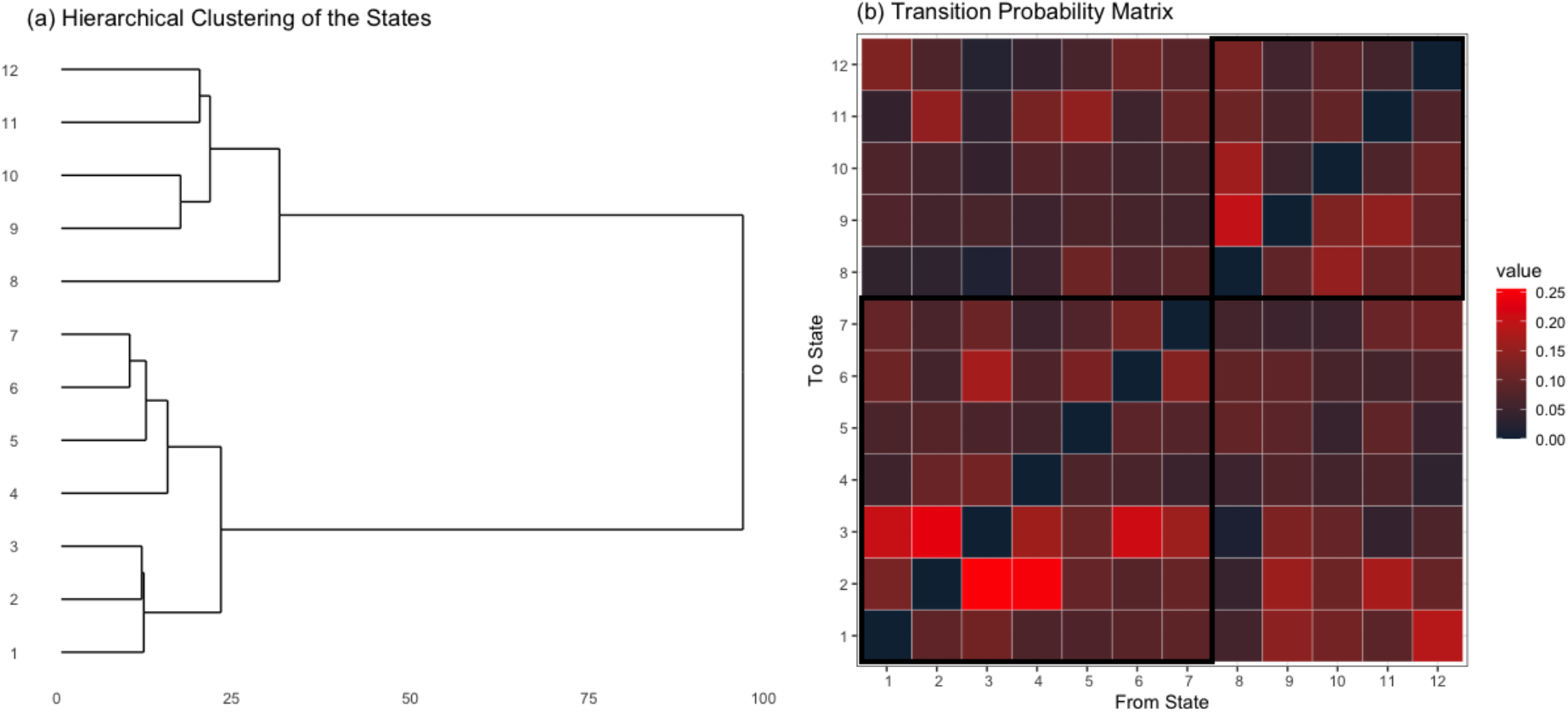
(a) Dendrogram for the covariance matrices across the 12 states. (b) Transition probability matrix for the approximate HSMM. States 1-7 and 8-12 correspond to metastates 1 and 2, respectively. These states are outlined in the black squares.

The above describes how we can approximate the HSMM with an HMM using state aggregates and a transition probability matrix with a specific structure. The use of state aggregates allows us to approximate dwell-time distributions within a state aggregate such that dwell-times are not necessarily geometric, and since the approximate HSMM is itself an HMM constructed in a special way we therefore inherit the theoretical and methodological properties and benefits of an HMM. Langrock and Zucchini (2011) developed the above strategy for univariate time series data, and we show using synthetic and empirical data that this strategy also works well for multivariate time series.

### 2.3 Estimation

Our use of an HMM to approximate an HSMM allows us to use the approaches developed for HMMs to estimate the parameters, being mindful of constraints, namely, the deliberate form of the transition probability matrix and the emission distribution parameters being the same within a state aggregate. In this work, we use the Expectation-Maximization (EM) algorithm (Dempster et al., 1977; Zucchini et al., 2017). We provide an overview of the algorithm here, adopting the notation by Zucchini et al. (2017).

If we know the true state sequences *S_t,n_* for each subject *n*, then we could optimize the complete likelihood given the observed data *y_1_,…, y_n_* and the known state sequences *S_t,1_,…, S_t,N_*. For *t* = 1,…, *T* and *n* = 1,…, *N*, let *u_i,n_(t)* = 1 if *S_t,n_ = i* and 0 otherwise, and for *t* = 2,…, *T* let *υ_i,j,n_(t)* = 1 if both *c_t–1,n_ = i* and *c_t–1,n_ = j* and 0 otherwise. Let 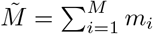 and let *b_ij_ = b_ij_(**β**)* denote the (*i,j*)-th entry of the matrix **B** = **B**(*β*), where the parameter ***β*** contains the parameters for the dwell-time distributions. Given the observed data and known state sequences, the complete data log-likelihood is

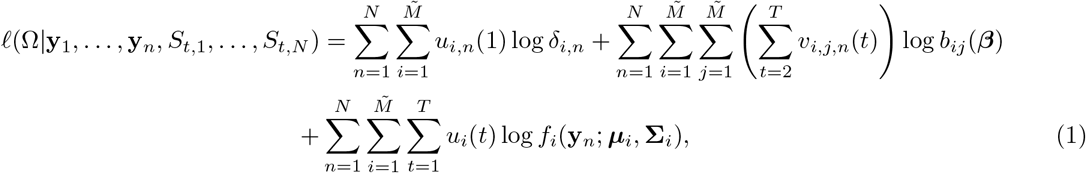

where Ω is the set of all parameters and *f_i_* is the probability density function for the *P*-variate normal distribution with mean vector ***μ**_i_* and covariance matrix **Σ**_*i*_. The EM algorithm iterates between the *E step* and the *M step* for optimizing Equation (1). The goal for the E step is to calculate the conditional expectation of {*u_i,n_(t)}_i,n_* and *{υ_i,j,n_(t)}_i,j,n_* given an estimate for Ω. On the other hand, the goal for the M step is to optimize Equation (1) replacing *{u_i,n_(t)}_i,n_* and *{υ_i,j,n_(t)}_i,j,n_* with their conditional expectations obtained from the E step. The EM algorithm alternates between these two steps until a convergence criterion is reached, which, in our case, is when there is a small relative change in the log-likelihood.

The E step remains the same as in standard HMMs, and so in the following we focus only on the M step. To update the transition probabilities in the M step we optimize the second term in Equation (1). There are two sets of parameters in this term: the parameter vector ***β*** = (***β**_1_*,…, ***β**_M_*) related to the dwell-time distribution and the transition probabilities **A** = (*a_ij_*). In this work for the dwell-time distribution we use a shifted Poisson distribution with shift parameter set to 1 and rate parameter *λ_i_*, which we relate to the covariates using the model log(*λ_i_*) = **Z*β_i_***, though other discrete distributions for the dwell-time distributions are possible. The shift parameter can be estimated, but we fixed that parameter here for simplicity. Thus, dwell-times potentially vary across states and could relate to covariates in different ways. To update ***β***, we numerically optimize the second term of Equation (1) using the `optim` function in `R`. Next, we use a closed-form solution to update **A** using *{υ_i,j,n_(t)}_i,j,n_* that was updated in the previous E step, namely, 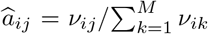, where 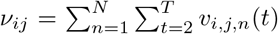. We point out that **B**(***β***) is a sparse matrix and so a number of its elements should be constrained to 0 and thus do not need to be optimized.

To update the parameters of the emission distribution in the M step we optimize the third term in Equation (1). A closed-form solution for these parameters exists. Since the parameters of the emission distribution must be the same across states within a state aggregate, then for state aggregate *i* = 1,…,*M*, we have

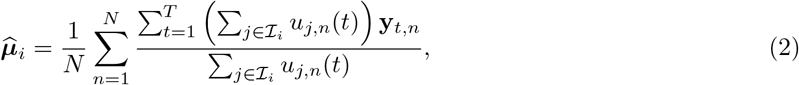

and

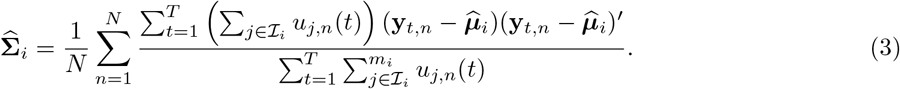

Recall that larger state aggregates improve the approximation to the dwell-time distribution but at a computational cost. Since the approximate HSMM is in the framework of an HMM, then the algorithmic complexity for the EM algorithm for the approximate HSMM is 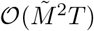 (Rabiner, 1989). In contrast, for a standard HSMM, the worst-case computational complexity for the EM algorithm is 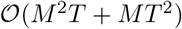 (Guédon, 2003). The computational complexity for the approximate HSMM remains linear with respect to the length of the time series even though we increase the dimensionality of the problem through the use of state aggregates. Larger state aggregates may be necessary depending on the context of the problem. In that case, one may want to take advantage of the sparse structure of the transition probability matrix **B**(***β***) to improve computational speed and for more efficient use of memory (Hadj-Amar et al., 2020).

## 3 Simulation Study

### 3.1 Simulation Settings

We conducted a number of simulations to show the performance of the approximate HSMM relative to other approaches under various conditions. In all cases, we simulated zero-mean *P*-variate time series data, *P ∈* {10, 30}, from a 3-state HSMM for *N* = 100 independent subjects, each time series having length *T* = 500. The emission distribution was a *P*-variate zero-mean normal distribution. The covariance matrices were as follows:

- **State 1. Σ**_1_ had a first-order autoregressive structure, such that its (*i, j*)-th element was set to 0.7^*|i-j|*^;
- **State 2. Σ**_2_ had a fourth-order autoregressive structure. The (*i, j*)-th element of the *precision matrix* (i.e., 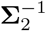) was set to **1**(*|i-j|* = 0) + 0.41(*|i-j|* = 1) + 0.21(*|i-j|* = 2) + 0.21(*|i-j|* = 3) + 0.11(*|i-j|* = 4), where **1**(·) is the indicator function that evaluates to 1 if its argument is true and 0 otherwise;
- **State 3. Σ**_3_ = (**Φ** + Id)^−1^, where Id is the *P × P* identity matrix and **Φ** is a random symmetric matrix such that an off-diagonal element was set to *1/P* with probability 0.5 or 0 with probability 0.5. Thus, the degree of sparsity remained constant as *P* varied, but the values of these elements were closer to 0 for larger *P*.

The above covariance matrices were used in the simulation study by Rothman et al. (2008) and Stadler and Mukherjee (2013), though we made slight modifications to **Σ**_3_. We simulated time series from an HSMM with transition probability matrix

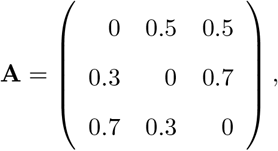

using a dwell-time distribution that was either i) a shifted Poisson distribution whose shift and rate parameters for the *i*-th state (1, *β_i_*), respectively, where log(***β***) = (2.5, 0.5, 1.5), or ii) a shifted negative binomial distribution whose shift, size, and mean parameters were (1, 10, *β_i_*), respectively, where ***β*** = (5, 10, 15). The purpose of the shift parameter is so that the dwell-time within a state is at least 1.

We assessed the performance of three competing models:

- **MHSMM.** We used the implementation of the HSMM in the `mhsmm` package in R (O’Connell et al., 2011). We set this model to have a Poisson distribution with rate parameter initialized to 1 for each state’s dwell-time distribution, transition probability matrix initialized to have 0.45 along the off-diagonal elements, and emission distribution parameters initialized to the maximum likelihood estimates obtained from segmenting each subject’s time series into three equal-sized segments.
- **HMM.** We fit an HMM using custom code. For the emission distribution, we initialized the mean vectors to the zero vector, and we initialized each state’s covariance matrix to have a compound symmetry structure by drawing the correlation from the Uniform (–*1/P, 1/P*) distribution and drawing the common standard deviation from the Uniform (1, 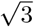) distribution. To ensure positive-definiteness of each state’s covariance matrix, we added 0.1 to the diagonal elements. We initialized the transition probability matrix to have 0.8 along the diagonals and 0.1 in the off-diagonals. We ran the EM algorithm and declared convergence when the relative change in the log-likelihood was < 10^−5^. We repeated this procedure 20 times, yielding 20 HMM fits, and we kept the fit that yielded the largest log-likelihood.
- **Approximate HSMM**. We fit the approximate HSMM described in Section 2. We set the size of the state aggregate corresponding to state *i* to *m_i_ = m ∈* {5, 10, 15}. We initialized the parameters of the emission distribution and the transition probability matrix to the estimates obtained by the HMM. We initialized the parameters of the dwell-time distribution (a shifted Poisson distribution with shift parameter 1 and rate parameter log(*λ_i_*) = *β_i_*) to a random draw from the normal distribution with mean 1.0 and standard deviation 0.5. Just as in the HMM, we used a tolerance criterion of 10^−5^ for the relative change in the log-likelihood for the EM algorithm, and after 20 random initializations, we kept the fit that yielded the largest log-likelihood.

After fitting each model, we addressed the label-switching problem by relabeling the states such that we minimize the Frobenius norm, denoted as ||·||_F_, between the estimated covariance matrices and the true covariance matrices across the three states. To assess model performance, we calculated the Frobenius norm between the estimated covariance matrix and the true covariance matrix for each state, and we report the estimate for 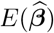. Note that whenever the true dwell-time distribution was the negative binomial distribution, both the MHSMM and the approximate HSMM were therefore deliberately misspecified since their dwell-time distributions were the Poisson distribution. Due to this misspecification, we instead compare 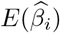 to the log of the center of dwell-time distribution for the *i*-th state. Finally, after relabeling the states, we reconstructed each subject’s state sequence using the Viterbi algorithm (Rabiner, 1989; Zucchini et al., 2017), and then calculated the misclassification rate averaged across time and subjects. We repeat each simulation study 100 times and averaged our assessments across the 100 simulations.

### 3.2 Simulation Results

In Table 1 we see how the approximate HSMM compares to the HMM and we also see the effects of different sizes of the state aggregates. The approximate HSMM for any state aggregate size has similar or better performances at estimating each state’s covariance matrix relative to the HMM. It may seem, however, that the approximate HSMM with state aggregate size *m* = 15 performed the worst at estimating the covariance matrix for State 1 in the scenario with *P* = 10 and shifted Poisson dwell-time distribution. Further investigation, however, is that there were some outliers that skewed the result. Indeed, the median error for the HMM was 0.134 and the median errors for the approximate HSMM was 0.061, 0.061, and 0.060 for *m* = 5, 10, and 15, respectively. It is not clear what led to the outliers in performance for *m* = 15. We used the shifted Poisson distribution as the dwell-time distribution, and thus it was correctly specified when the true dwell-time distribution was the shifted Poisson distribution and misspecified when it was the shifted negative binomial distribution. Thus, 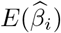 is an estimate of the mean of the dwell-time distribution for state *i*. When the true dwell-time distribution was the shifted Poisson distribution, the approximate HSMM generally estimated the true ***β*** well except for State 1, where we see a slight downward bias when *m* = 10 and a larger downward bias when *m* = 5. For State 1, the dwell-time distribution was truly centered at exp(2.5), and so the state aggregates were too small, similar to the example we showed in Figure 5. We draw similar conclusions when the true dwell-time distribution was the shifted negative binomial distribution. Note that due to the misspecification of the dwell-time distribution, we compare 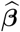 to the log of the true means, (log(5), log(10), log(15)). As before, we see the approximate HSMM underestimating the true mean of the dwell-time distribution for States 2 and 3 for *m* = 5 and State 3 for *m* = 10. While larger values of *m* led to a reduction in bias in estimating the mean of the dwell-time distribution, we also see that larger values of *m* also led to greater variability in the estimates. Finally, we see that the misclassification rate for the approximate HSMM were comparable over the different values of *m*. In all cases, the approximate HSMM for any state aggregate size had a similar or better performance than the HMM with respect to misclassification. The low misclassification rate for all methods under all scenarios is likely because of how distinct the covariance matrices were across the three states.

**Table 1:**
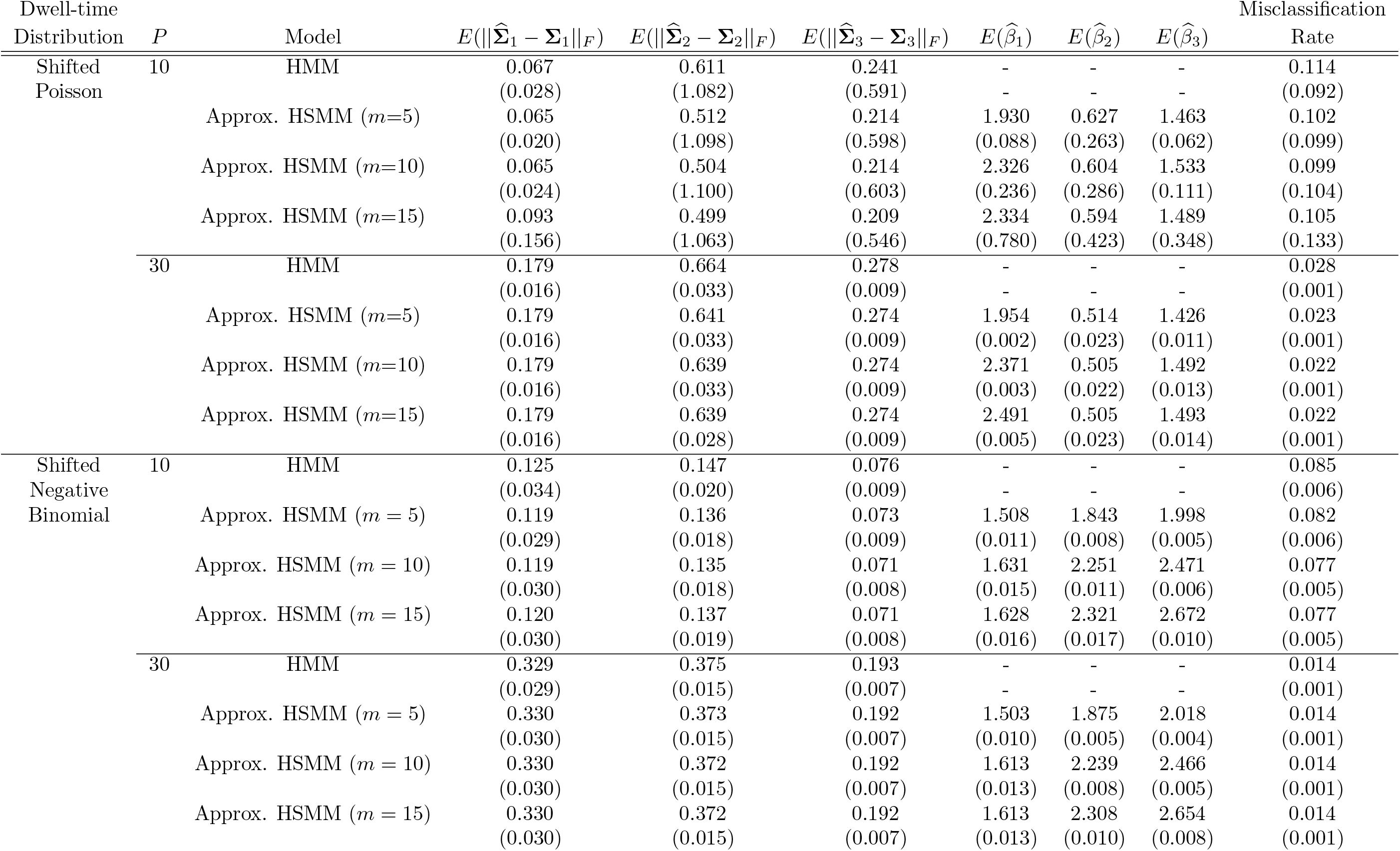
Simulation results showing the performance under various conditions of the HMM and the approximate HSMM with different sizes, *m*, for the state aggregates, reported as the average (SD) across the 100 simulations. The HMM does not have a parameterized dwell-time distribution, hence its entries corresponding to ***β*** are marked with a hyphen (-).

**Figure 5:**
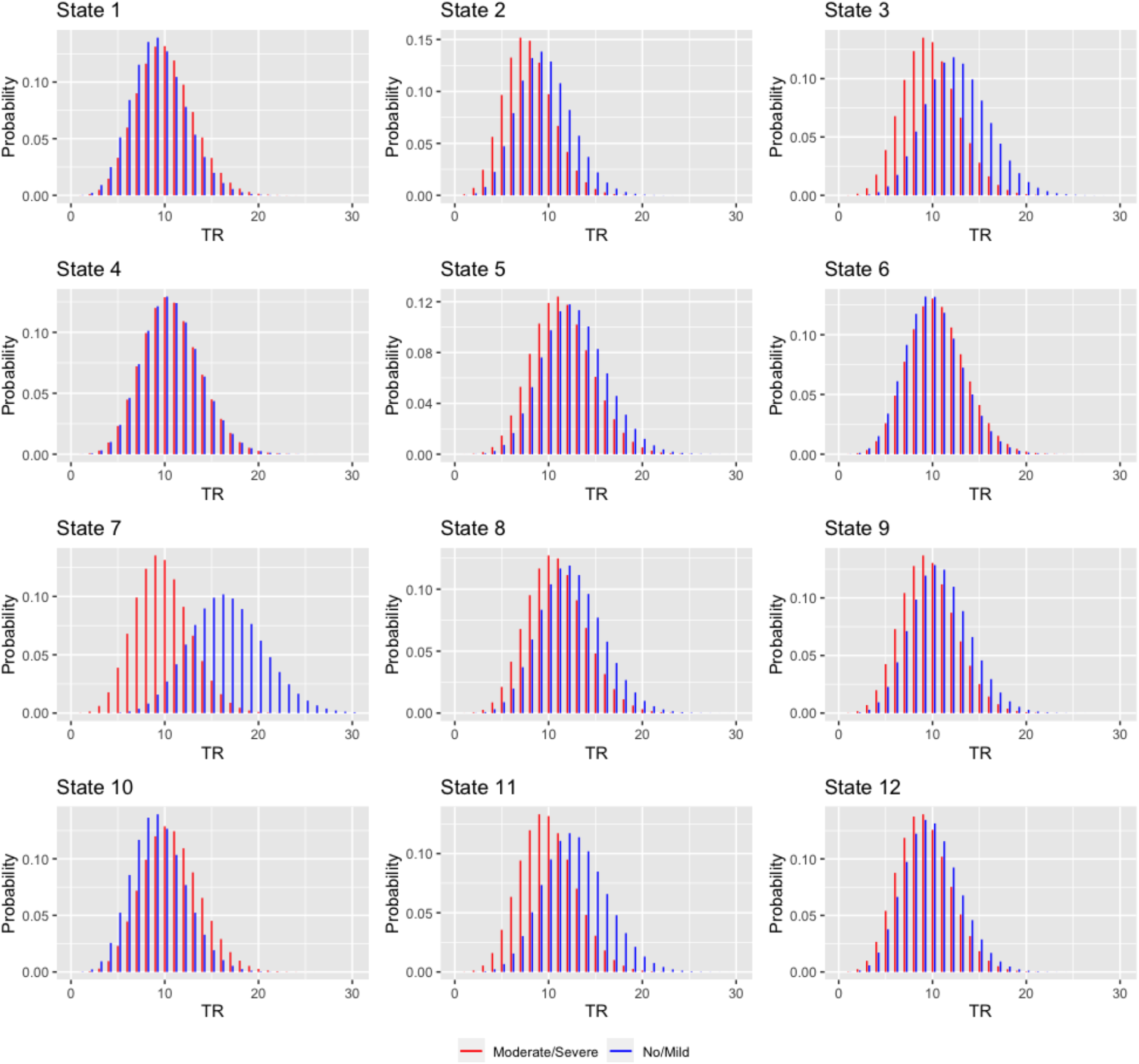
Dwell-time distributions for each state, stratified by NSSI severity group.

In Table 2 we compare the performances of the MHSMM, the HMM, and the approximate HSMM. First, we point out that in some instances for MHSMM, the estimates of ***β*** diverged to −∞. We removed these instances when tabulating results, and so the results for MHSMM reported here are more optimistic than they truly are. With respect to estimating each state’s covariance matrix, as we saw before generally the approximate HSMM performed as well as or improved on the HMM. We also see that the MHSMM and the approximate HSMM have similar performances in estimating each state’s covariance matrix. However, when it comes to estimating the mean of the dwell-time distributions, we see that the MHSMM has mixed results. Indeed, the MHSMM had good performances on aveage, but the estimates could have higher variance or could diverge, as stated earlier. Finally, the misclassification performance of the HMM and the approximate HSMM were either as good as or better than the misclassification performance of the MHSMM.

**Table 2:**
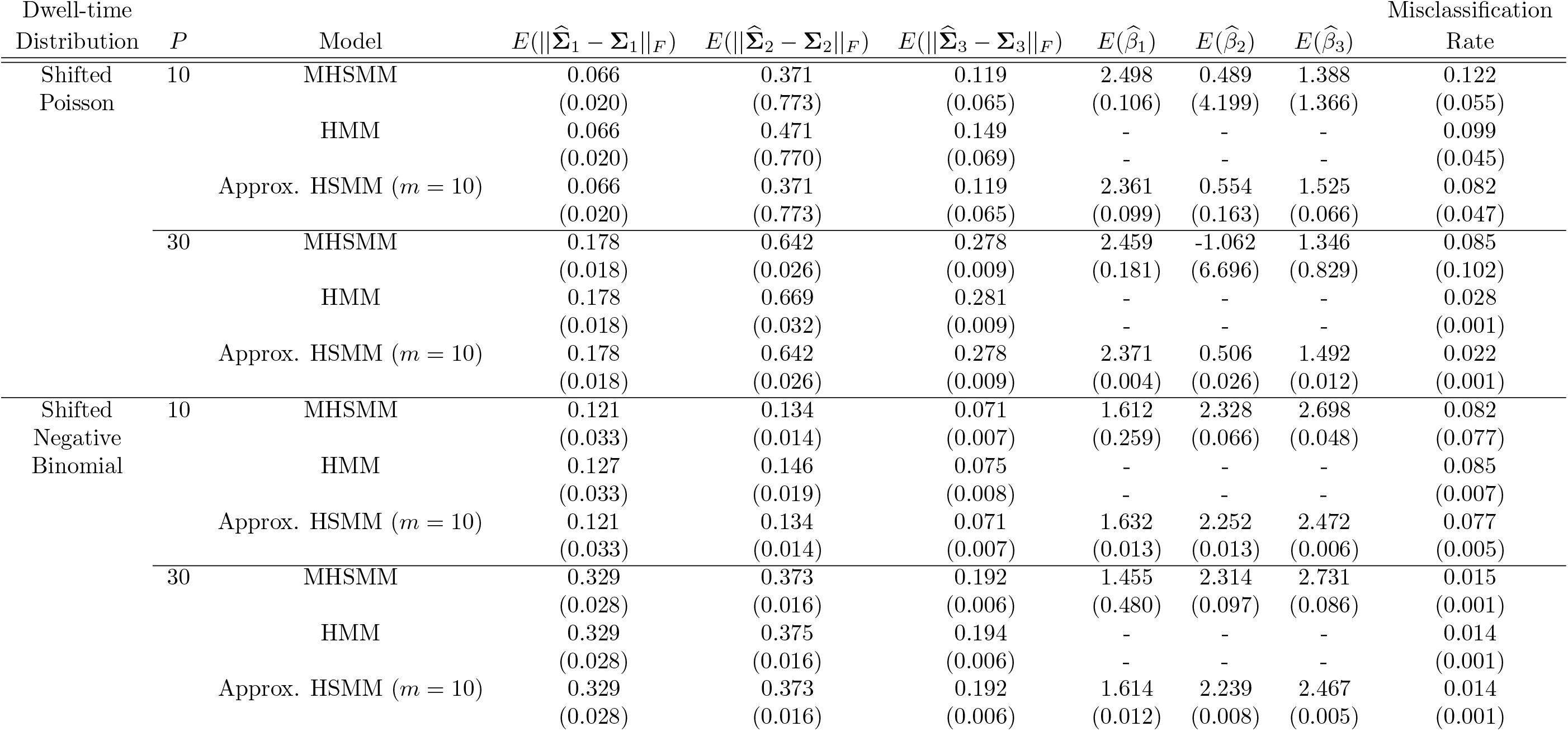
Simulation results showing the performance under various conditions of the MSHMM, the HMM, and the approximate HSMM *m* = 10 reported as the average (SD) across the 100 simulations. The HMM does not have a parameterized dwell-time distribution, hence its entries corresponding to ***β*** are marked with a hyphen (-).

In summary, our simulation study shows that the approximate HSMM performs well at estimating the covariance matrices, and its performance at estimating the mean of the dwell-time distribution depends on the state aggregates. We initialized the emission distribution parameters of the approximate HSMM to the estimates of the emission distribution parameters of the HMM, which explains why the approximate HSMM improved on the HMM. We believe that this is good practice given how how fast one can fit the HMM. Finally, while the approximate HSMM seems to have greater bias than the MHSMM in estimating the mean of the dwell-time distribution, the estimates of the MHSMM can be relatively unstable.

## 4 Applications to Dynamic Connectivity Analysis

### 4.1 Data Description

The data came from the BRain Imaging Development of Girls’ Emotion and Self (BRIDGES) Study at the University of Minnesota. A central goal of the BRIDGES Study is to understand the how the development of brain systems may be aberrant in adolescents with with non-suicidal self-injury (NSSI). This longitudinal study recruited adolescent females 12-16 years of age who exhibited a continuum of NSSI severity which was then classified into 4 categories of NSSI severity: no NSSI, mild NSSI, moderate NSSI, or severe NSSI. The adolescents were enrolled to participate in three annual evaluations that involved clinical, physiologic and neuroimaging assessments. The current study focused on the baseline neuroimaging assessment, and considered two clinical groups for comparison: adolescents with moderate or severe NSSI versus those with no or mild NSSI.

Brain scanning sessions were conducted at the Center for Magnetic Resonance Research at the University of Minnesota. The resting-state functional magnetic resonance imaging (fMRI) data consisted of a 12-minute scan during which each participant was instructed to stay awake, keep their eyes open focused on a fixation cross, and to “not think about anything in particular”. These fMRI scans consisted of whole brain T2*-weighted functional volumes with 2 mm isotropic voxel resolution, with the following fMRI parameters: TR = 800ms, TE = 37 ms, flip angle = 52°, FOV = 212 mm, 2 mm isotropic voxel, Multiband factor=8. All functional data were acquired using the Human Connectome Project multiband echo planar imaging sequence.

Group level spatial ICA was utilized to estimate intrinsic connectivity networks (ICNs) using the GIFT toolbox (https://trendscenter.org/software/gift/) (Calhoun et al., 2001). Subject level time courses were reduced to 110 principal components and concatenated along the time dimension. The concatenated time courses were reduced to 100 principal components through a group level PCA. Group level ICN’s were estimated using the infomax algorithm to optimize temporal independence (Calhoun et al., 2001). Calculations on group level ICN’s were repeated 20 times using the ICASSO technique for reliability (Himberg and Hyvarinen, 2003). The spatial maps from the infomax ICA output were used as spatial templates for Group Information Guided ICA (GIG-ICA). The GIG-ICA calculations estimated 100 group level ICN’s also optimizing the independence at the subject level (Du and Fan, 2013; Du et al., 2020). 35 ICN spatial maps were selected to be binarized, setting the bottom 45% nonzero absolute voxel intensities to zero, and evaluated for mutual overlap with ROIs from the Yeo 17 network and the Harvard-Oxford cortical and subcortical structural atlases (https://identifiers.org/neurovault.collection:262) (Yeo et al., 2011). Each ICN was labeled using the atlas region with which it had the most overlap.

The final data used in our analysis was a time series with length *T* = 912, dimension *P* = 35, observed from *N* = 100 subjects, and with each time series having zero mean and unit variance. Of these 100 subjects, 61 were classified to have moderate or severe nonsuicidal self-injury (NSSI), and 39 were classified to have no or mild NSSI.

### 4.2 Overview of the Statistical Procedure

Vidaurre et al. (2017), Vidaurre et al. (2018), and Shappell et al. (2019) each fit a 12-state model to their resting-state fMRI data, and so we selected the same model to reproduce their results. We set the covariate matrix **Z** to have two columns, where the first column is an indicator for moderate or severe NSSI and the second column is an indicator for no or mild NSSI. For the dwell-time distribution for the *i*-th state we used a shifted Poisson distribution with shift parameter set to 1 and rate parameter log(*λ_i_*) = **Z*β_i_***, where 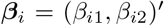 captures the relationship between NSSI severity and dwell-time. For the *i*-th state, we set m_i_ = 20. Thus, the approximate HSMM had a total of 240 states.

We now describe how we fit the approximate HSMM. As in our simulation study, we first fit a standard HMM with 12 states using the EM algorithm. We used the same strategy as in our simulation study for the random initializations, so we refer to Section 3 for details. We ran the EM algorithm and declared convergence when the relative change in the log-likelihood was < 10^−6^. After obtaining fits from 50 random initializations, we kept the fit that yielded the largest log-likelihood.

We used the estimates of the parameters of the emission distribution for the HMM and its transition probability matrix (setting its diagonal entries to 0 and then standardizing the off-diagonals to have a unit row sum) to initialize the parameters for the EM algorithm to fit the approximate HSMM. To initialize the dwell-time distribution ***β***, we randomly drew each coefficient from the normal distribution with mean 2.5 and standard deviation 0.5, so that the resulting prior dwell-time for each state would be roughly between 1 and 32.5 TR units (or between 0.8 and 26 seconds). We ran the EM algorithm and declared convergence when the relative change in the log-likelihood was < 10^−5^. We repeated this procedure 50 times, yielding 50 approximate HSMM fits, and we kept the fit that yielded the largest log-likelihood. We then used the `numDeriv` package in R to numerically estimate the Hessian matrix for ***β*** to obtain standard errors and to construct Wald test statistics for post-hoc analyses about these coefficients. We used the Viterbi algorithm for state reconstruction for each subject (Zucchini et al., 2017), and then mapped each state sequence over the 240 states back to the 12 state aggregates. Given each subject’s state reconstruction, we computed their state switching frequency, defined as a transition from one state aggregate to another, and their fractional occupancy, defined as the proportion of time spent in a state. Finally, to assess the similarity between the 12 state aggregates, we used hierarchical clustering, using the Kullback-Liebler divergence between covariance matrices as a measure of distance between states.

### 4.3 Results

Henceforth, in the context of the approximate HSMM we will use the terms “state” and “state aggregate” interchangeably. Figure 3 shows the correlation matrix for each of the 12 states in the approximate HSMM. We see across all 12 states that that the different time series within a region generally maintained strong correlations. However, across the 12 states we see varying degrees of the strength of the correlation between regions. For instance, States 8, 9, and 12 are characterized by strong inter-region correlations, whereas States 3 and 7 are characterized by weak inter-region correlations.

Figure 4(a) shows the similarity between the 12 states. We see that the 12 states can be separated into two *metastates*, in agreement with previous findings (Vidaurre et al., 2017; Shappell et al., 2019), with States 1-7 form Metastate 1, and States 8-12 form Metastate 2. Vidaurre et al. (2017) and Shappell et al. (2019) showed that transitions from one state to another are greater if the two states are within the same metastate. Our results are in partial agreement. Figure 4(b) shows the transition probability matrix. States 2 and 3 tend to transition to each other. State transitions between the states in metastate 1 tend to be to other states within metastate 1, specifically to States 2 or 3, but we see that states within metastate 2 tend to transition to States 1-3 which are in metastate 1.

Table 3 shows the estimates of ***β*** across all states, which we then mapped back to the rate parameter of the shifted Poisson distribution as shown in Figure 5. The dwell-time distributions were generally centered around TR = 10, corresponding to 8 seconds. Looking at Figure 5, we see that in many of the states the Moderate/Severe group tended to have shorter dwell-times. To test if the dwell-time distributions differed between NSSI severity groups, we tested *H_0_: β_i2_* – *β_i1_* = 0 for each *i*, then used the Benjamini-Hochberg procedure to correct for the false discovery rate (FDR) over the 12 comparisons (Benjamini and Hochberg, 1995). The difference was statistically significant (FDR-adjusted *p*-value < 0.05) for all states except for State 4.

**Table 3:** Parameter estimate (standard error) for the dwell-time distribution for each state and each NSSI severity group. We controlled the *p*-values for the 12 comparisons using the false discovery rate procedure.

| State | NSSI Severity | | Adjusted<br>$p$-value |
| --- | --- | --- | --- |
|  | No/Mild | Moderate/Severe |  |
| State 1 | 2.107 (0.019) | 2.201 (0.014) | < 0.001 |
| State 2 | 2.126 (0.017) | 1.926 (0.015) | < 0.001 |
| State 3 | 2.437 (0.018) | 2.169 (0.015) | < 0.001 |
| State 4 | 2.261 (0.021) | 2.268 (0.016) | 0.788 |
| State 5 | 2.445 (0.020) | 2.344 (0.016) | < 0.001 |
| State 6 | 2.196 (0.018) | 2.248 (0.016) | 0.035 |
| State 7 | 2.737 (0.018) | 2.168 (0.014) | < 0.001 |
| State 8 | 2.418 (0.016) | 2.284 (0.015) | < 0.001 |
| State 9 | 2.271 (0.016) | 2.149 (0.013) | < 0.001 |
| State 10 | 2.101 (0.015) | 2.268 (0.012) | < 0.001 |
| State 11 | 2.455 (0.016) | 2.186 (0.015) | < 0.001 |
| State 12 | 2.174 (0.021) | 2.094 (0.017) | 0.003 |

As discussed above, the shorter dwell-times for those with Moderate/Severe NSSI suggest a greater number of state switches over the course of the resting-state scan. In Figures 6(a) and 6(b), we see this to be the case. The mean (SD) number of state switches is 102.69 (9.01) in the Moderate/Severe NSSI severity group and 94.08 (9.49) in the No/Mild NSSI severity group (*p*-value < 0.001 using a two-sample *t*-test). The mean (SD) number of metastate switches is 56.07 (8.00) in the Moderate/Severe NSSI severity group and 49.21 (8.67) in the No/Mild NSSI severity group. Table 4 shows the fractional occupancy for each of the 12 states and each of the 2 metastates. We see that States 2 and 3 were the most visited states for both groups, with the remaining 10 states being visited somewhat equally. However, the dwell-time parameters shown in Table 3 do not suggest that States 2 and 3 have longer dwell-times relative to the other states. Thus, the larger fractional occupancy is due to the number of transitions into States 2 and 3 as opposed to the longer dwell-times. There is no evidence of a difference in fractional occupancy between the NSSI severity groups for any state or metastate (*p*-value > 0.05 for all comparisons).

**Figure 6:**
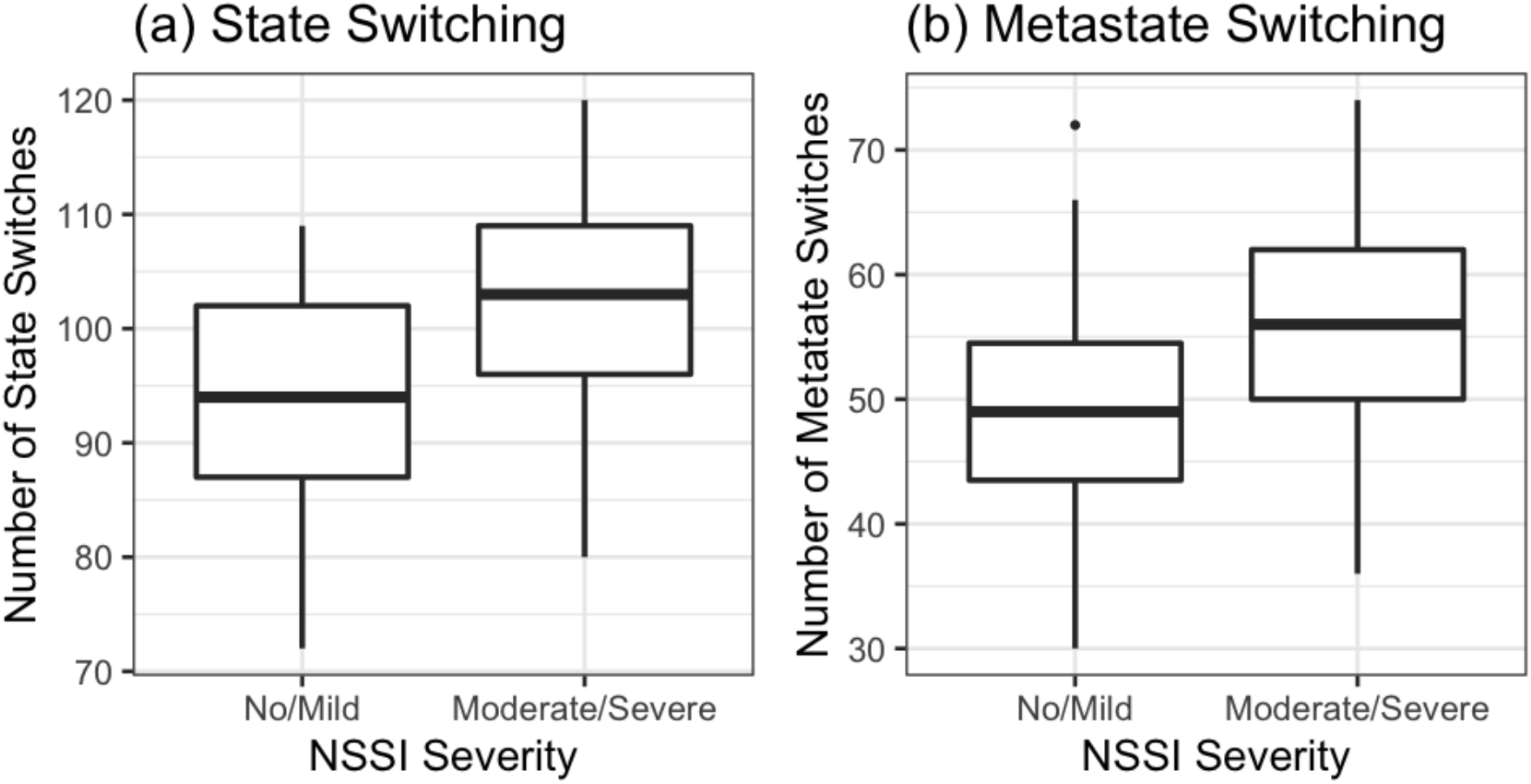
Boxplots of (a) state-switching frequency and (b) metastate-switching frequency per NSSI severity group.

**Table 4:** Mean (SD) fractional occupancy per state and metastate for each NSSI severity group.

| State | Fractional Occupancy |  |
| --- | --- | --- |
|  | No/Mild | Moderate/Severe |
| 1 | 0.075 (0.034) | 0.088 (0.042) |
| 2 | 0.105 (0.050) | 0.109 (0.056) |
| 3 | 0.124 (0.095) | 0.137 (0.076) |
| 4 | 0.069 (0.034) | 0.071 (0.032) |
| 5 | 0.083 (0.037) | 0.071 (0.036) |
| 6 | 0.093 (0.049) | 0.100 (0.049) |
| 7 | 0.079 (0.050) | 0.083 (0.034) |
| 8 | 0.080 (0.080) | 0.065 (0.066) |
| 9 | 0.081 (0.033) | 0.076 (0.034) |
| 10 | 0.063 (0.046) | 0.060 (0.039) |
| 11 | 0.086 (0.047) | 0.076 (0.044) |
| 12 | 0.061 (0.028) | 0.063 (0.033) |
| Metastate 1 | 0.544 (0.097) | 0.531 (0.090) |
| Metastate 2 | 0.456 (0.097) | 0.469 (0.090) |

## 5 Discussion

We developed a statistical model for approximating HSMMs using an HMM, and showed its utility in analyzing resting-state fMRI time series. Our analysis showed that dwell-time distributions vary over states, and differ between NSSI severity. Our results are consistent with previous work on evidence of two metastates, but we did not see strong evidence that states tended to transition to other states within the same metastate (Vidaurre et al., 2017; Shappell et al., 2019). It may be the case that the transition probabilities between states is associated with NSSI severity which we did not account for in our model. Indeed, in our analysis NSSI severity was only included in the dwell-time distributions. Previous work has shown that the strength of connectivity and the variability in the temporal dynamics of connectivity between specific regions of the brain can be useful for characterizing depression and suicide ideation (Demirtaş et al., 2016; Kaiser et al., 2016; Jiao et al., 2019; Wise et al., 2017). It is possible for our approximate HSMM model to focus on the dynamic connectivity between specific region pairs by fitting a collection of bivariate models. Our findings show that female adolescents with moderate or severe NSSI severity have more state switches, suggesting greater temporal variability in the correlations. Although no prior studies have examined dynamic connectivity in relation to NSSI, a small number of prior studies utilized different techniques to study resting-state connectivity in patients with depression. Different directions of association between temporal variability and connectivity have been previously reported, but these studies used different analytic approaches (e.g., sliding-window correlations, cluster analysis, graph theory), and time series data from different regions of the brain (Demirtaş et al., 2016; Kaiser et al., 2016; Jiao et al., 2019; Wise et al., 2017; Zheng et al., 2018; Zhi et al., 2018). Notably, these prior studies have been primarily conducted on adults, and no prior studies have applied HMMs or HSMMs to examine state duration and transition frequency in depressed patients. Furthermore, while depression and NSSI are related constructs in the sense that NSSI usually occurs in the context of negative affect (Klonsky, 2011) and the two problems commonly co-occur, growing evidence suggests that depression and NSSI might have both overlapping and distinct underlying biological patterns (Klimes-Dougan et al., 2019). Finally, we showed that fractional occupancy was the greatest for State 3. We point out that State 3 is characterized by weak correlations averaged across all the off-diagonal entries of its correlation matrix (mean correlation = 0.173), with only State 7 having weaker correlations (mean correlation = 0.145), although Marusak et al. (2017) showed that healthy children spend more time in a state characterized by weak regional connections.

Our approximate HSMM has natural extensions that can provide the practitioner with further flexibility. In our application, we accounted for the NSSI Severity group in the dwell-time distribution, but our use of the HMM framework means we could model the parameters of the emission distribution or in the transition probability between state aggregates (Zucchini et al., 2017). However, we may encounter challenges with respect to updating the parameters in the M-step of the EM algorithm. Furthermore, we restricted our analysis to *P* = 35 dimensions. One could consider a model with higher dimensions, though one may need to include some form of regularization in the model. While there have been theoretical and methodological developments for HMMs for high-dimensional time series (Fiecas et al., 2017b; Städler and Mukherjee, 2013), to our knowledge there have been no developments for HSMMs. Since our approximate HSMM is in the HMM framework, one could therefore utilize the developments for HMMs for modeling high-dimensional time series data. Another potential extension is to account for the autocorrelation of the data, which could improve estimates of and statistical inference on functional connectivity (Fiecas et al., 2017a; Hart et al., 2018). One possibility is to use a switching vector autoregressive (VAR) model, which uses a stationary VAR conditional on the state sequence (Ting et al., 2017; Ombao et al., 2018). Finally, our analysis only used the baseline data. One could conduct a longitudinal dynamic connectivity analyses by incorporating random effects into the model. There have been developments for HMMs with random effects (Altman, 2007; DeRuiter et al., 2017), and one could potentially adapt these developments to the approximate HSMM.

We now describe some limitations with our modeling approach. In our analyses, we assumed that the number of states was known *a priori*. This will not always be the case in practice, and so one would need to use, e.g., the Akaike or Bayesian information criterion (AIC or BIC, respectively) or other model comparison metrics to select the number of states (Zucchini et al., 2017). In our approximate HSMM, we also have the size of the state aggregates as a tuning parameter, and we showed that this can affect performance if it is too small with respect to the true underlying dwell-time distribution. In our data analysis, we set the size *m_i_* = 20 for all state aggregates and so our approximation of dwell-time distributions will be accurate for up to 16 seconds. We saw in our analysis that dwell times were generally centered at *TR* = 10, corresponding to 8 seconds, and thus the right-tail of the dwell-time distribution was likely not affected by our choice for *m_i_*. Indeed, the flexibility of the HSMM means that there one would need to make more decisions about the model and tune more parameters. There are a number of open problems and questions surrounding dynamic connectivity (Lurie et al., 2020). For instance, it is important to demonstrate if the summary measures extracted from our approximate HSMM (e.g., state-specific correlations, state reconstructions) are robust and reproducible. We showed the performance of the approximate HSMM using synthetic data, but empirical evidence will be more useful. The literature is mixed on the test-retest reliability of summary measures extracted from dynamic connectivity analyses (Choe et al., 2017; Liao et al., 2017). On the other hand, Vidaurre et al. (2018) showed that HMMs yielded reproducible results in modeling brain dynamics. Finally, it is important to be clear on what we mean when we say that connectivity is “dynamic”, since this can influence the appropriate “null model” to use to test for the presence of such dynamics and can also affect the interpretation of results (Liégeois et al., 2017; Lurie et al., 2020). In our case, the approximate HSMM, like the HMM, assumes that the correlations vary over time conditional on the state sequence. However, we point out that unconditional on the state sequence the first and second-order moments do not vary over time.

Altogether, we showed using synthetic data and resting-state fMRI data that the approximate HSMM provides an excellent modeling framework for conducting dynamic connectivity analyses in resting-state fMRI. The theoretical and methodological foundations of the approximate HSMM are based on those already established for HMMs, and thus this framework has the flexibility to be extended or adapted based on the needs of the practitioner. Finally, the model is fairly general, and even though we used this model to analyze resting-state fMRI data, it can be adapted to analyze multivariate time series data from other scientific fields.

## Notes

### Competing Interest Statement

The authors have declared no competing interest.

